# The architecture and operating mechanism of a cnidarian stinging organelle

**DOI:** 10.1101/2021.09.09.459669

**Authors:** Ahmet Karabulut, Melainia McClain, Boris Rubinstein, Sean A. McKinney, Matthew C. Gibson

## Abstract

The stingers of jellyfish, sea anemones and other cnidarians, known as nematocysts, are remarkable cellular weapons used for both predation and defense^1^. Nematocysts are specialized organelles which consist of a pressurized capsule containing a coiled harpoon-like thread^2^. These structures are in turn built within specialized cells known as nematocytes^3^. When triggered^4^, the capsule explosively discharges, ejecting the coiled thread which punctures the target and rapidly elongates by turning inside out in a process called eversion^5,6^. Due to the structural complexity of the thread and the extreme speed of discharge, the precise mechanics of nematocyst firing have remained elusive^7^. Here, using a combination of live and super-resolution imaging, 3D electron microscopy and genetic perturbations, we define the step-by-step sequence of nematocyst operation in the model sea anemone *Nematostella vectensis*. This analysis reveals the complex biomechanical transformations underpinning the operating mechanism of nematocysts, one of the nature’s most exquisite biological micro-machines. Further, this study will provide insight into the form and function of related cnidarian organelles and serve as a template for the design of bioinspired microdevices.

## Introduction

Cnidarian nematocysts are complex cellular weapons with highly specialized forms and functions^2^. Nematocysts are Golgi-apparatus derived intracellular organelles comprised of venomous threads enclosed within a pressurized capsule^8^. When triggered, the capsule discharges, ejecting its thread as a harpoon which penetrates targets, delivering a cocktail of neurotoxins^9,10^. At the cellular level, nematocyst discharge is among the fastest mechanical processes in nature, known to be completed within 3 milliseconds in *Hydra* nematocysts^11,12^. This high-speed discharge is generated by the accumulation of osmotic pressure inside the capsule by a matrix of cation binding poly-γ-glutamate polymers (PGs)^13,14^. During its subsequent operation, the nematocyst thread undergoes a shape transformation, turning inside-out through a process called eversion which is caused by the release of both osmotically generated pressure and elastic energy stored in the thread^15,16^.

Nematocyst characteristics vary significantly among different cnidarian species, exhibiting diversity in capsule size and thread morphology, but all retain a similar mechanism of operation^17–19^. We studied the operation of the nematocyst thread in the genetic model sea anemone *Nematostella vectensis* which harbors two types of nematocysts: microbasic p-mastigophores and basitrichous isorhizas, the later having short and long varieties^20,21^. In sea anemones, nematocyst capsules are sealed by three apical flaps connected to the stinging thread^22^. This thread is composed of two distinct sub-structures: a short, rigid and fibrous shaft and a long thin tubule decorated with barbs^16,18^. The shaft is composed of three helically coiled filaments, and is initially ejected as a compressed projectile, piercing the target, and later everts to form a lumen through which the remainder of the thread, the tubule, is released^16^.

While it is known that shaft eversion entails a geometric transformation from a tightly compressed coil to a hollow syringe, the mechanisms driving this process are poorly understood. Further, tubule eversion significantly differs from that of the shaft, as the tubule everts by simply turning inside-out^23^. The release of the capsule pressure is sufficient to drive the initial ejection and penetration of the shaft, however, additional energy sources are likely to be required for further elongation of the tubule^7,15,24^. Due to the speed and complexity of these events, the precise stages of discharge and eversion have thus far remained elusive. Here, we investigate the structural composition and mechanical transformations of both the shaft and the tubule during distinct phases of nematocyst discharge in *Nematostella vectensis*.

### Visualizing the nematocysts of *N. vectensis*

To understand the distribution of stinging cells (nematocytes) and their nematocysts in *Nematostella*, we first created a transgenic line expressing EGFP in nematocytes under the control of the *nematogalectin* promoter region (*nematogalectin*>*EGFP*; Fig. 1a). Live imaging of transgenic primary polyps showed that the tentacles were predominantly populated with nematocytes bearing the long form basitrichs (Fig.1 a^I^). The body column was populated with the shorter variety along with a few p-mastigophores. Intriguingly, we found that nematocytes were connected through neurite-like processes which formed local networks (Fig. 1a^II^, *arrow*). Nematocytes are known to form synapses and act as afferents or effectors but can also operate cell-autonomously^25–28^. Thus, the observed networks might function in regulating collective behavior and coordinated activity of nematocyte populations^29^. In *EGFP*+ nematocytes, fluorescence was detected throughout the cytoplasm and the sensory apparatus but was excluded from the capsule (Fig. 1b). The capsule wall and thread are built, in part, of minicollagens which allow construction of a variety of structural fibers by cross-linking^30–33^. We exploited this to visualize the capsule content by treating live animals with fluorescent TRITC which was incorporated into the nematocyst thread during its maturation, presumably through a reaction with minicollagens^34,35^. We were therefore able to analyze the architecture of the thread from its development to its final morphology after firing (Fig. 1b, c; Supplementary Video 1). In contrast to the dense shaft of p-mastigophores (Fig. 1c, *arrows*), in which the dye intensity was very high compared to the tubule, fluorescent TRITC incorporated with similar intensity in both the shaft and tubule of basitrichs (Fig. 1c, *dashed arrows*). The more uniform labeling of basitrichs and their prevalence in primary polyps led us to investigate the thread operation in this nematocyst type.

**Fig. 1.**
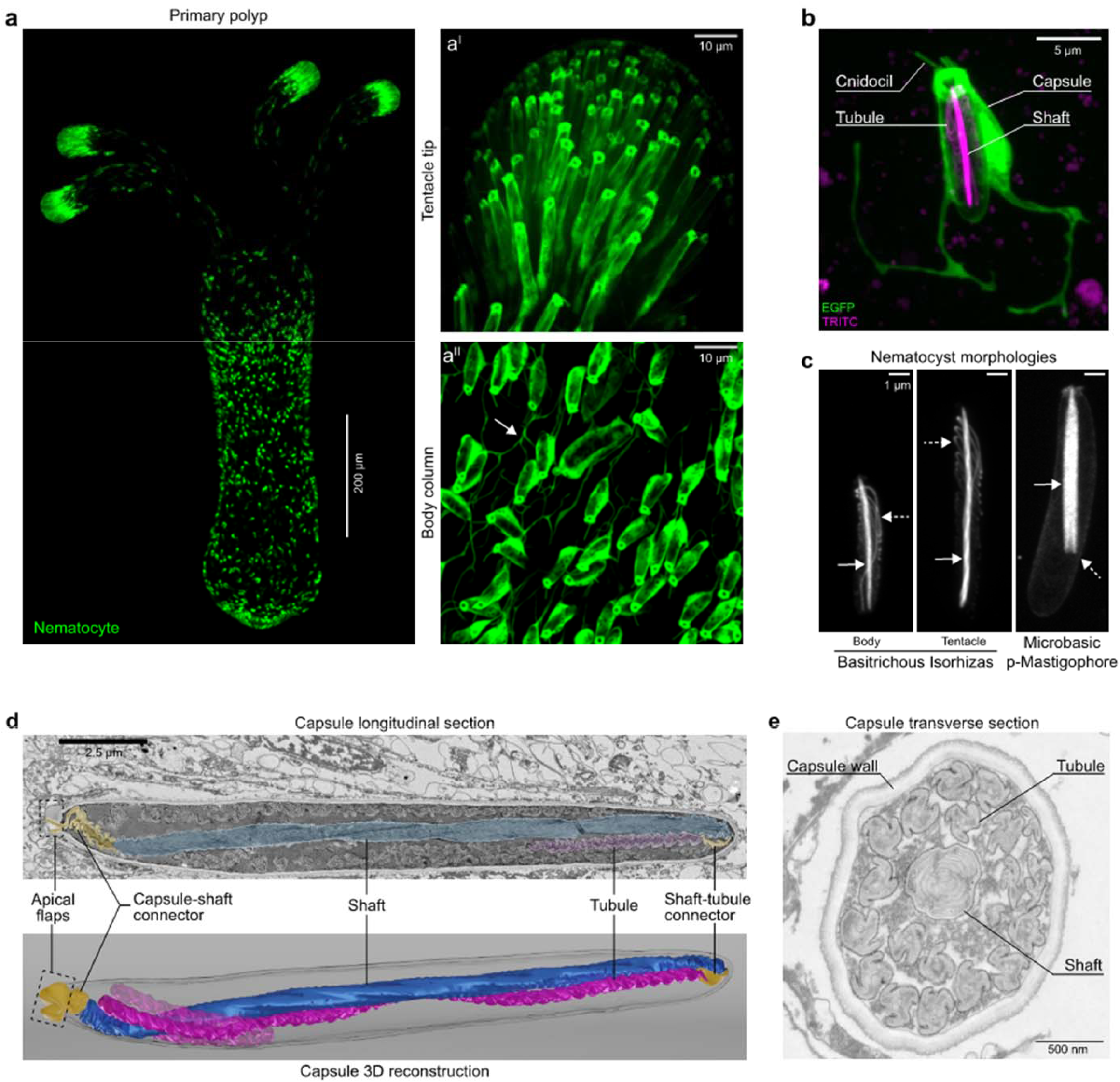
The architecture of undischarged nematocysts. **a,** Nematocytes (*green*) of a transgenic *N. vectensis* primary polyp expressing EGFP under the control of the *nematogalectin* promoter. **a^I^,** EGFP expression at the tentacle tip. **a^II^,** EGFP expression on the body column. The arrow points to the network of cellular processes connecting nematocytes. **b,** Close-up view of a single nematocyte on the body column of a primary polyp expressing EGFP in nematocytes (green). The cnidocil (sensor) apparatus, cell body and neurite-like processes are shown. TRITC (magenta) labels the capsule, the centrally positioned shaft and the folds of the tubule. **c,** Nematocyst morphologies in *N. vectensis* based on the fluorescence signal of incorporated TRITC. Basitrichous isorhizas with densely labeled shaft (arrow) and coiled tubule (dashed arrow) continuous with the compressed shaft (left and middle panels). Microbasic p-mastigophores shaft (arrow) with its distinctive V-shaped notch (right panel, dashed arrow). Scale bars 1 μm. **d,** Longitudinal section of a nematocyst showing densely coiled shaft filaments (blue), a portion of the coiled tubule (magenta) and the two connector regions, the capsule-shaft connector and shaft-tubule connector (yellow). The apical flaps are seen in a partially open conformation (dashed box). Corresponding 3D reconstruction of the longitudinal section shows the capsule, central shaft (blue), a portion of the attached tubule (magenta), connector regions (yellow) and apical flaps (dashed box). **e,** Transverse section of a nematocyst showing the capsule wall, dense lamellar shaft and the propeller shaped tubule.

### The architecture of undischarged nematocysts

To analyze the structure of the shaft and tubule and thereby determine their functionality, we next performed 3D-reconstruction of undischarged basitrich capsules from serial sections using scanning electron microscopy (SEM; Fig. 1d; Supplemental Video 2). We found that the compressed shaft consisted of tightly coiled filaments vertically aligned to the capsule aperture formed by the apical flaps (Fig. 1d, *box*). The filaments were composite structures with stacks of lamellae built from electron-dense and electron-lucent layers. The thread wall encasing these filaments was connected to the apical flaps with a loose capsule-to-shaft connector. A similar shaft-to-tubule connector was located between the basal end of the shaft and the apical end of the tubule. The tubule was twisted, forming pleats in regular lengthwise segments (Supplemental Video 2, 3). In capsule cross sections, the shaft lamellae were observed to be tightly coiled and compressed, while the tubule cross section exhibited a propeller-shaped structure (Fig.1e; Supplementary Video 4).

### Nematocyst discharge and eversion of the stinging thread

To determine the distinct phases of thread operation, we recorded fluorescent high-speed video of discharge events in TRITC-treated animals (Supplementary Videos 5-7). Following *in situ* nematocyte stimulation, the compressed shaft was first ejected as a dense projectile which then rapidly expanded to form an elongated cylinder through which the tubule emerged (Fig 2a^II^-a^IV^, *arrows;* Supplementary Video 5). Based on these observations, we defined three principal phases of nematocyst operation: shaft discharge (Phase I), shaft eversion (Phase II) and tubule eversion (Phase III; Fig. 2, *box*). Using SEM, we visualized the ultrastructure of the discharging shaft. In the undischarged state, we observed sparse lamellae decorating the region where the shaft tapered to the capsule-shaft connector (Fig. 2b, *dashed arrow*). During the early stages of discharge, the everted capsule-shaft connector formed a skirt around the traversing shaft, creating a double-walled structure (Fig. 2c, *arrow*) with the uneverted shaft moving forward inside the connector (Fig. 2c, *blue*). The everted capsule-shaft connector was externally covered with sparse filaments resembling irregular spines originating from the eversion of the lamellae observed in the undischarged state (Fig. 2c, *dashed arrow*). Serial SEM sections also captured an everted connector (Fig. 2d, *arrow*) in which the tubule could be observed departing the capsule (Fig. 2d *dashed arrow;* Supplementary Video 8). Finally, in SEM sections of a partially discharged nematocyst thread, we observed the uneverted tubule traversing inside of its everted fraction, which was decorated externally with hollow barbs (Fig. 2e, *arrow*; Supplementary Video 9).

**Fig. 2.**
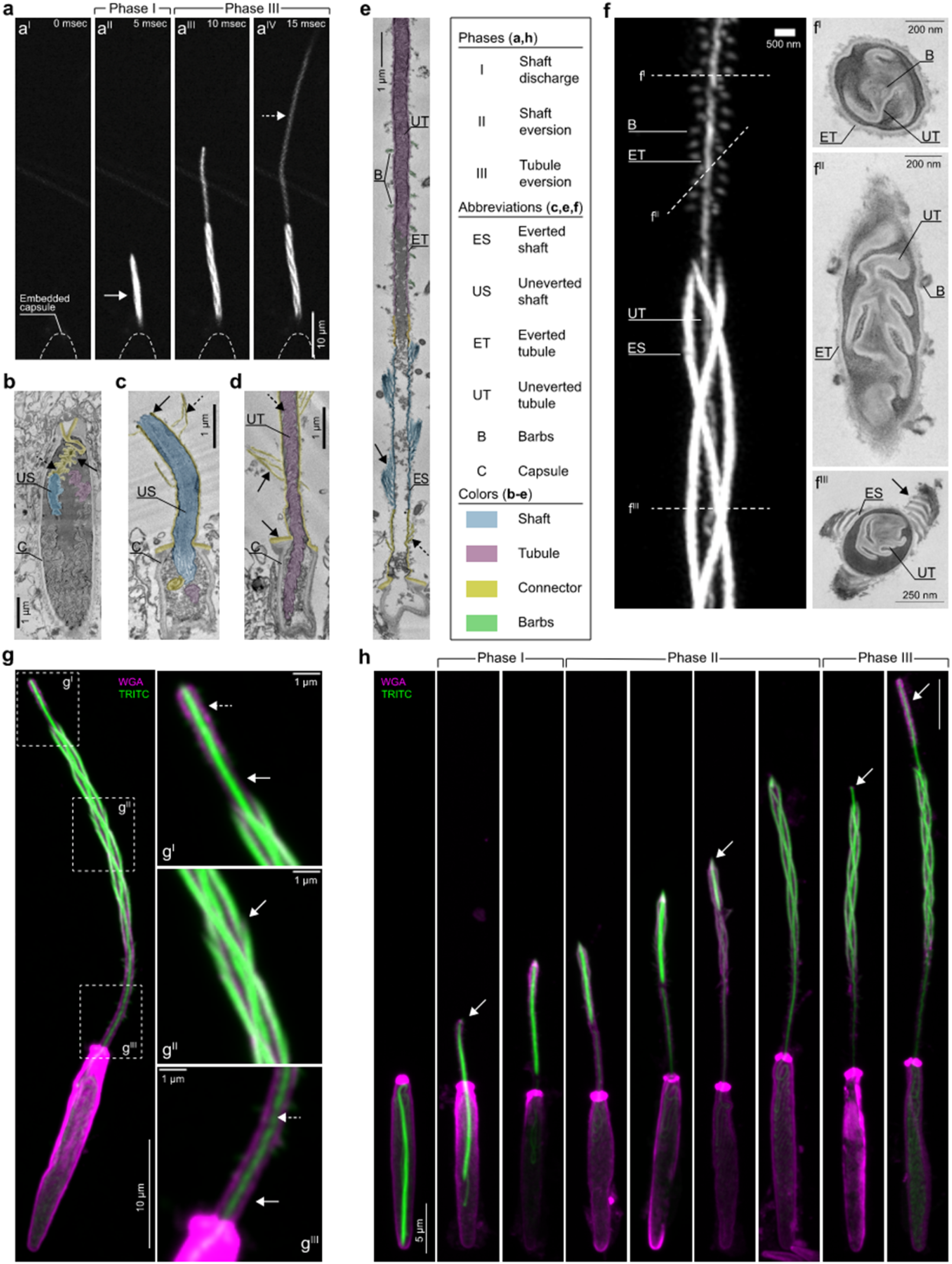
Nematocyst discharge and the mechanism of shaft eversion. **a,** Live imaging of nematocyst discharge. TRITC labeled nematocysts were discharged *in vivo* and frames were captured at 5 milliseconds intervals. **a^I^**-**a^IV^,** Snapshots showing distinct phases corresponding to the shaft discharge and tubule elongation. Note: The shaft eversion (Phase II) was too fast to be captured in this sequence. **b,** Longitudinal section of a capsule (C) prior to discharge showing the uneverted shaft (US) and its taper (dashed arrow) to the shaft-tubule connector (yellow, arrow), the apical flaps (V-shaped), and the uneverted tubule (magenta). **c,** Longitudinal section of a discharged capsule during shaft eversion showing shaft filaments (US, blue), capsule-shaft connector (outside the capsule, yellow), a portion of the tubule (magenta) and shaft-tubule connector (yellow).The double-walled structure (arrow) and sparse lamellae on the connector’s exterior (dashed arrow) are also shown. **d,** Longitudinal section of an everted shaft-capsule connector (yellow, arrows) and traversing uneverted tubule (UT, magenta, dashed arrow). **e,** Longitudinal section of a partially everted tubule. The everted shaft (ES, blue, arrow), capsule-shaft connector (yellow, dashed arrow), parts of the uneverted tubule (UT, magenta) inside the everted tubule (ET) and barbs (B) are shown. **f,** A partially everted nematocyst thread revealed by TRITC incorporation. The image shows the everted shaft (ES) filaments, the uneverted tubule (UT, center) and the barbs (B) decorating the everted portion of the tubule (ET). Corresponding EM cross sections: **f^I^,** Cross section of the tubule. Labels indicate the everted and uneverted tubule segments with barbs (B) located at the center. **f^II^,** Oblique section of the partially everted tubule showing tubule segments and barbs at the exterior (B). **f^III^,** Cross section of the everted shaft (ES) and the traversing uneverted tubule (UT). The electron-dense and -lucent layers of the ES filaments (*arrow*) are also shown. **g,** A partially discharged nematocyst stained with fluorescent dye-conjugated wheat germ agglutinin (WGA, magenta) and TRITC (green), showing the shaft fibers (green) and the thread wall (magenta). Magnified regions: **g^I^,** The shaft-tubule connector (arrow) and the everting tubule emerging with barbs (dashed arrow). **g^II^,** The shaft filaments (green, arrow) and the WGA labeled thread wall (magenta). **g^III^**, The capsule-shaft connector (arrow) showing the thread wall (WGA, magenta) and the traversing uneverted tubule (TRITC, green, dashed arrow). **h,** Representative images of the distinct phases of thread eversion. Fluorescent staining shows the geometric transformations in each phase. Arrows indicate the apex of the thread during distinct phases.

To better understand the structural changes described above, we next analyzed super-resolution images of TRITC-labeled threads undergoing eversion. This approach allowed us to demonstrate the existence of a triple helical geometry of the uncoiled shaft filaments together with the traversing uneverted tubule which could be traced by the labeling of the barbs (Fig. 2f). Interestingly, the thread wall did not incorporate TRITC and was invisible in fluorescent images but could be seen in corresponding SEM cross-sections as an electron-lucent layer that enclosed the uneverted tubule in its compacted state (Fig. 2f^I^, 2f^II^). The highly ordered arrangement of barbs within the uneverted tubule indicates that these structures are stacked as a column, which appeared as a single filament in fluorescent images (Fig. 2f, 2f^I^). Further, SEM cross sections through the shaft showed that its filaments consisted of lamellae which enclosed the traversing uneverted tubule, as seen in fluorescent images (Fig. 2f, 2f^III^, *arrow*).

Next, to visualize the thread wall that was otherwise invisible in optical images, we focused our attention on non-collagenous components of the thread. Together with minicollagens, the nematocysts of *Hydra* and *Nematostella* contain glycans and show similarities to the extracellular matrix in composition^8,36–38^. A nematocyst-specific lectin, Nematogalectin, acts as a scaffold linking the minicollagens to glycans, mainly consisting of a non-sulfated chondroitin sheath^39,40^. Considering the presence of lectins in the structure, we hypothesized that the presence of GAGs could be detected with fluorescently labeled sugar-binding lectins. Thus, we stained discharged nematocysts with fluorescent dye-conjugated Wheat Germ Agglutinin (WGA), which is selective for GlcNAc chains, and found that WGA strongly bound to the electron-lucent thread wall (Fig. 2g)^41^. Co-staining with TRITC showed that the WGA-stained material did not incorporate TRITC, but rather formed a laminate with TRITC-labeled structures. Finally, images of threads in an early everted state showed that the TRITC-labeled shaft surrounded the WGA-labeled tubule wall. The connector regions lacking filaments or barbs which were poorly labeled with TRITC also became visible with WGA staining (Fig. 2g^I^-g^III^). In conclusion, these results suggest that the thread consists of two layers: a collagen-rich layer forming the TRITC-detectable shaft filaments and barbs, and a GAG enriched layer forming the overall cylindrical thread wall, including the connector regions, shaft wall and tubule wall.

### The mechanism of shaft eversion

Structural studies of nematocyst threads penetrating gel substrates indicate that shaft eversion is initiated at the shaft’s apex^16^.To determine how the shaft transforms from its compressed state to a loose triple helical structure, we captured the early stages of discharge by treating *Nematostella* primary polyps with a solution that simultaneously triggers discharge and rapidly fixes the samples^20^. A reconstructed sequence of events from still images revealed the complex geometric transformation of the shaft as it exited the capsule in a coiled configuration (Fig. 2h). We found that during shaft discharge (Phase I), the ejected shaft continued to move forward as a dense projectile inside the capsule-shaft connector until the connector extended to its maximal length. During shaft eversion (Phase II), the filaments started to uncoil from the apex of the shaft, turning inside out, thereby everting, while the basal end of the shaft moved forward inside the uncoiling filaments. The tubule was attached to the basal end of the shaft through the shaft-tubule connector and was thus pulled through the newly formed lumen inside the uncoiled shaft filaments. This movement resulted in complete eversion of the three filaments where the shaft’s former apical end became its basal end. Finally, the everted shaft lumen opened, permitting the movement of the shaft-tubule connector, initiating Phase III. The emergence of barbs on the everted tubule exterior demarcated a boundary between the shaft-tubule connector and the tubule itself, and thus marked the beginning of tubule eversion (Fig. 2g^I^; Fig. 2h, last panel). Altogether, these data indicate that shaft eversion executes a reproducible series of physical transformations that involve the uncoiling and forward motion of its filaments.

### The mechanism of tubule eversion

SEM and fluorescent images revealed differences in composition and structure between the triple-helical fibrous shaft and the smooth cylindrical tubule (Fig. 3a, b). While shaft eversion can be explained by the motion of three filaments, the tubule lacks such geometry and likely everts by a mechanism that involves the unfolding and untwisting of the tubule wall^15,42^. Live imaging revealed that during tubule elongation the forward-moving tubule untwisted and relaxed to a cylindrical state (Fig 2a^III^-a^IV^; Fig 3b, *Stages 1*, *2*; Supplementary Video 5). At the everting tip, the helical twists of the everting tubule segment could be seen due to the concentration of WGA staining along the barb pockets (Fig. 3c). In contrast, in an image captured shortly after the initiation of tubule eversion, the tubule was seen as a double-walled cylindrical structure with a lumen between the uneverted and everted walls, suggesting that it likely relaxed and untwisted rapidly (Fig. 3d, *arrows*). These results indicate that tubule eversion likely occurs in stages involving untwisting of its propeller-like shape to a cylindrical conformation, and that the action of the everted segment untwisting and relaxing feeds forward the remaining uneverted tubule to the distal tip. Co-staining with the *Nematostella* minicollagen Ncol4 antibody also revealed an overlap with the TRITC signal, indicating that the barbs are made of minicollagens, and TRITC likely labels, in part, minicollagen rich fibers (Fig. 3e).

**Fig. 3.**
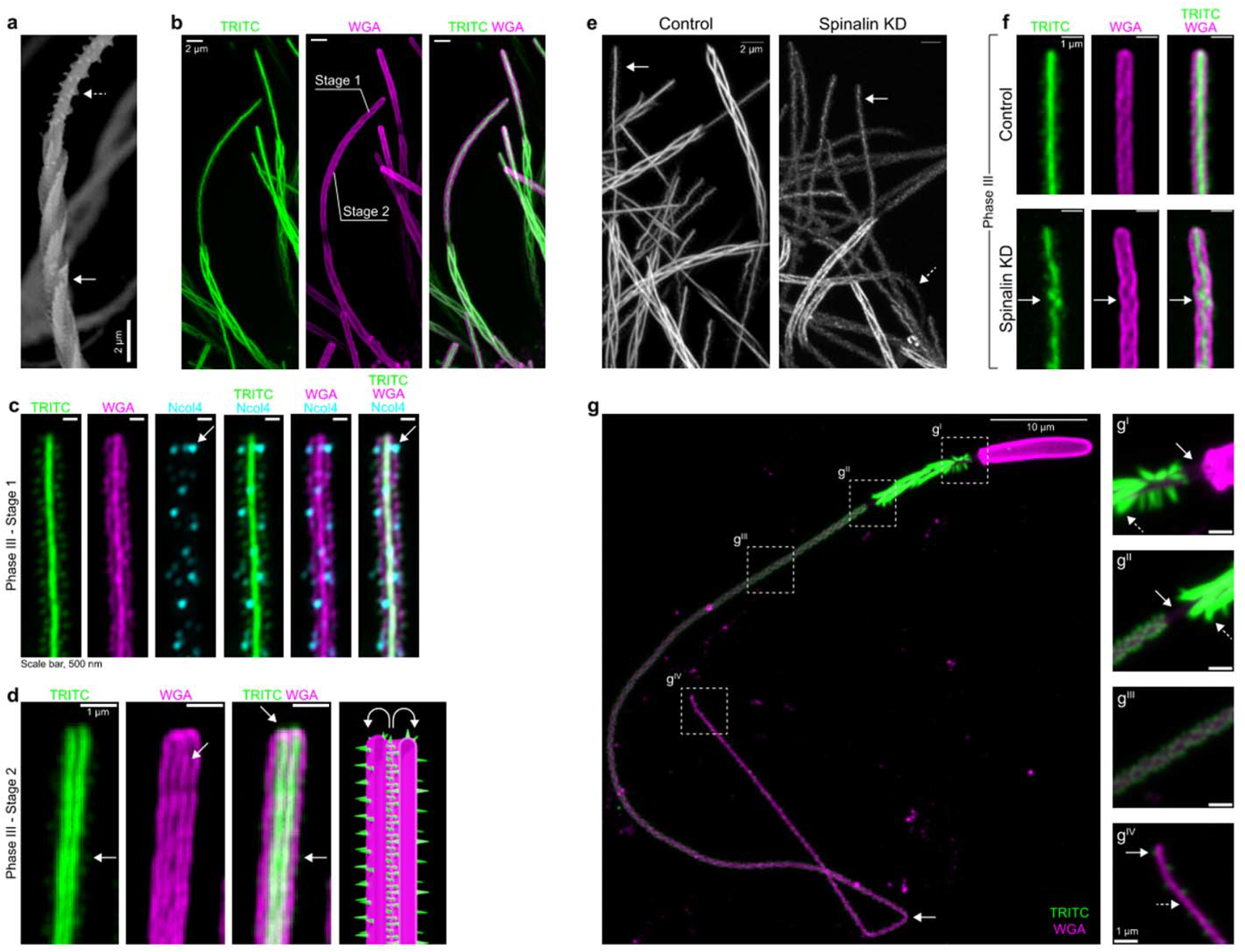
The mechanism of tubule eversion. **a,** SEM image of the shaft (arrow), tubule and helically arranged barbs (dashed arrow). **b,** Fluorescent image of a partially everted tubule. The shaft filaments and barbs (TRITC, green) and the thread wall (WGA, magenta) are shown. Scale bars 2 μm. **c,** Super-resolution image of the everting tubule tip (Stage 1). The tubule (WGA, magenta), barbs (TRITC, green) and co-labeling with anti-NCol4 antibody (*cyan*, *arrow*) are shown. Scale bars 0.5 um. **d,** The tip of an everting tubule (Stage 2). The tubule wall (magenta), its double-walled structure (left middle panel, arrow) and the barbs (right middle panel, arrow) are shown. Illustration of Stage 2 (right panel). Scale bars 1 μm. **e,** Structure of the tubule barbs (arrows) in control and *spinalin* KD samples (dashed arrow). Scale bars 2 μm. **f,** Effects of the spinalin knockdown on the barbs (TRITC, green) and tubule wall (WGA, magenta). The arrows indicate the disorganized barbs. Scale bars 1 μm. **g,** A fully everted thread labeled with TRITC and WGA. Arrow indicates kinking at a distal tubule site. **g^I^,** Capsule-shaft connector (arrow) and the everted shaft (green, dashed arrow). **g^II^**, The shaft-tubule connector (arrow) and the everted shaft (dashed arrow). **g^III^**, Fully everted tubule wall (magenta) and the barbs (green). **g^IV^**, Tip of the thread showing the tubule wall (magenta, dashed arrow) sparsely decorated with barbs (green, arrow).

To test the role of the centrally stacked barbs in tubule eversion, we use shRNA^43,44^ to knock down the reported *Nematostella spinalin* gene (*v1g243188*)^45^, which encodes a fibroin-like protein and possible ortholog of *Hydra Spinalin*^46^. This glycine- and histidine-rich protein was reported to be present in the spine structures on the surface of everted tubules in *Hydra* nematocysts^47^. In *Nematostella*, we found that knockdown resulted in weakly labeled, thinner shaft filaments and disrupted the structure of the barbs. The loss of TRITC intensity indicated that, like minicollagens, a fraction of the dye was also incorporated into material enriched with this protein. While the knockdown disrupted the structure of the barbs and their arrangement (Fig. 3e, *arrows*), this did not appear to affect thread operation. However, loss of *spinalin* resulted in increased bending of the tubule compared to controls (Fig. 3e, *dashed arrow*). This observation suggests that *spinalin* is a major component of the thread which is required for structural integrity of the thread but not its operation. Further, the stereotypical helical arrangement of the barbs and their stacked configurations were disrupted (Fig. 3f, *arrow*). In discharged nematocysts, the fully everted thread appeared to be an isodiametric tube composed of a WGA-labeled wall equipped with TRITC-labeled barbs and shaft filaments, excluding the connector regions (Fig. 3g, *dashed boxes*). The barbs decreased in density from the proximal to the distal end of the tubule, and sparsely decorated the distal region (Fig. 3g^II^-g^IV^). We hypothesize that the barbs, internally stacked before eversion and externally helically distributed after eversion, might function as a skeleton that prevents further bending and kinking for the elongating tubule. Indeed, as the barbs lessened distally, the tubule appeared to become more prone to kinking compared to the proximal regions which could bend in smooth curves (Fig. 3g, *arrow*). Interestingly, in live-imaging of an elongating thread, the tubule performed 180° turns in its barb dense proximal region, likely following the path of least resistance (Supplementary Video 10). Altogether, our data indicate that tubule eversion involves the unfolding of the tubule wall in which barbs likely provide structural support for the elongating thread.

### Model of the geometric transformation of the shaft and tubule

These findings allowed us to build a model describing the key aspects of the observed geometric eversion in three phases. Our results suggest that the shaft filaments are attached apically to the capsule flaps, and basally to the tubule via connector regions (Fig. 4a). **Phase I**: Upon discharge, the shaft is ejected along with the capsule-shaft connector covering the ejecting shaft. A double-walled structure is formed (Fig. 4b,c). Based on still images and videos, shaft eversion occurs after complete ejection of the shaft. Thus, we postulate that the connector accumulates maximal elastic stress when the ejected shaft reaches its maximal distance from the capsule (Fig. 4c). **Phase II**: Elastic stress on the capsule-shaft connector creates outward forces applied to the apex of the compressed shaft filaments resulting in detachment of the filaments and initiation of the eversion process due to the release of elastic stress within the shaft (Fig. 4d-g, *initiation*). Upon completion of the sequence, a lumen is formed within the shaft that is protected by the thick filaments (Fig. 4h, *end of Phase II*). **Phase III**: The final phase commences with the release of the shaft-tubule connector which everts by folding on itself forming a double-walled structure (Fig. 4i). The uneverted segment of the tubule then progressively exits the capsule and moves through the everted portion of the elongating thread (Fig. 4j).

**Fig. 4.**
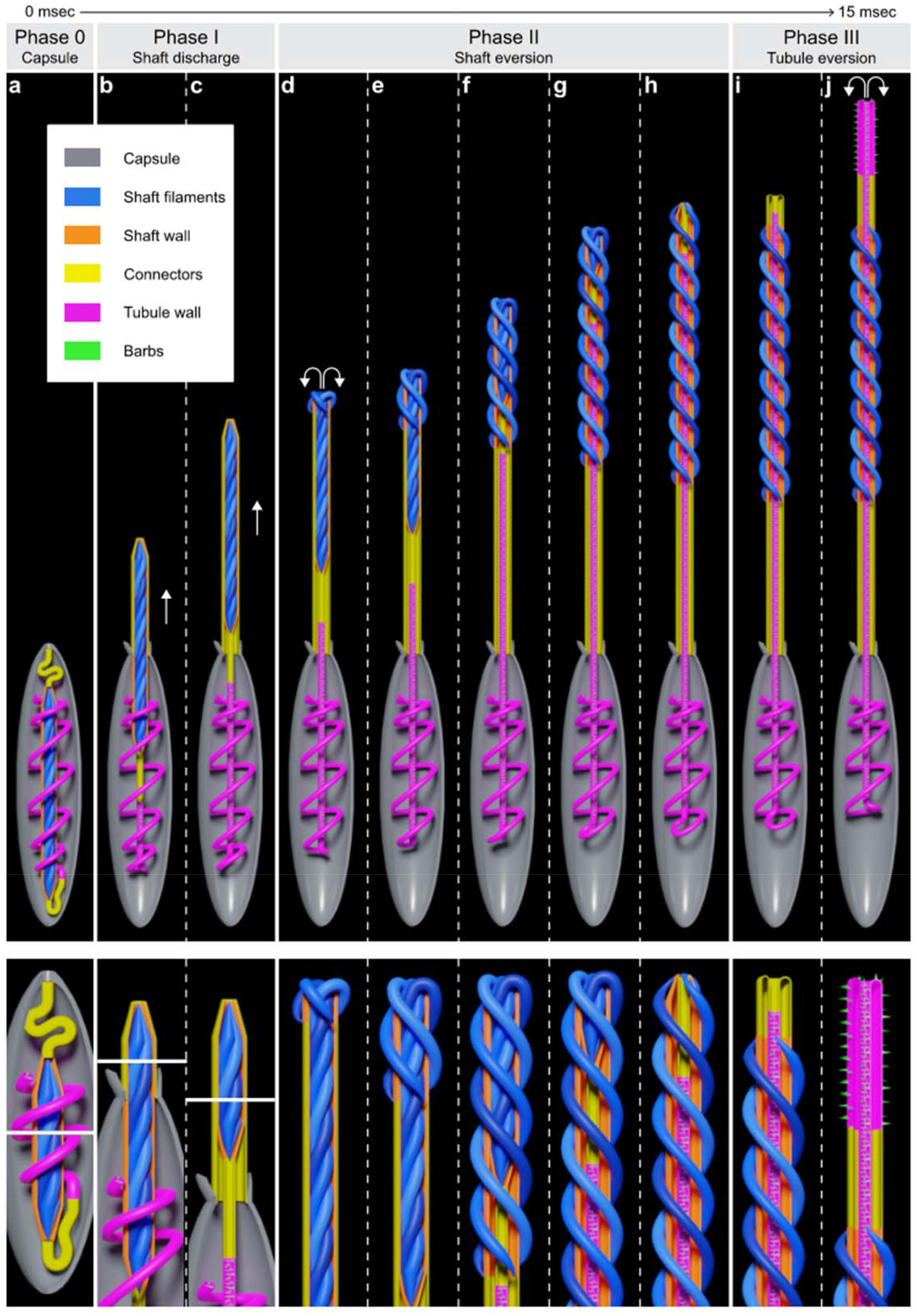
Model of the geometric transformation of the shaft and tubule. The box indicates the sub-structures of the nematocyst. Lower panels: Magnified views of critical regions during distinct phases. **a,** Undischarged capsule with tightly coiled shaft surrounded by the shaft wall, two connectors, and the coiled tubule. **b-c,** Initial stage of shaft discharge (Phase I). The forward movement and eversion of the capsule-shaft (CS) connector enclosing the ejected shaft filaments are shown. The arrows indicate the forward movement of the shaft. **d-h,** The geometric eversion mechanism of the shaft and uncoiling of the compressed shaft filaments (Phase II). Arrows indicate the direction of the forces applied to the apex of the compressed shaft filaments. **e-g,** Steps in the progression of shaft eversion. The model shows the uncoiling shaft filaments and the forward movement of the shaft-tubule connector and the tubule. **h,** The final stage of shaft eversion. Note: The basal end of the uneverted shaft becomes the apical end of the everted shaft. **i-j,** The eversion mechanism of the shaft-tubule connector and the tubule (Phase 3).

## Discussion

In this article, we described the 3D organization of the nematocyst and the sequence of geometric transformations that occur upon its activation. We also suggest a model explaining the specific mechanisms of thread eversion. Based on our results, we conclude that nematocyst operation occurs in three stages involving a complex transformation of the shaft and the elongation of the tubule, during which energy stored in the overall structure is transformed to kinetic energy. The shaft performs two critical functions: first as a compressed syringe to penetrate the target cuticle; second as a protective tunnel for passage of the thin tubule. The shaft eversion process resembles the mechanics of a Y-shaped slingshot wherein elastic energy is stored in two bands attached to a pad containing a projectile. Upon release of the pad, the stretched bands experience a geometric eversion. The elastic energy stored in the bands converts into kinetic energy of the accelerating projectile. Nematocysts utilize a similar approach, but due to the limited space inside the capsule store the elastic energy required for shaft eversion by bending and twisting three filaments and attaching them to the capsule and the tubule. The helices of the everted filaments exhibit a larger radius and step compared to their uneverted configuration. Thus, the amount of elastic energy stored in the helical coils decreases in the everted state. The excess energy released during shaft eversion is possibly used for the release of the shaft-tubule connector (Supplementary Information). The connectors might function in the initiation of eversion; stretching of the capsule-shaft connector utilizes energy from the discharge to initiate shaft eversion, while the shaft-tubule connector transfers the elastic energy of the “slingshot” to initiate tubule eversion. The loose shaft-tubule connector rapidly forms a cylindrical tube which might act as a buffer zone for the transition of the three-filament shaft to the twisted tubule.

The source of the driving force for tubule penetration could be explained by the osmotic pressure and the elastic forces accumulated in the tubule structure. It has been shown that hydration of ruptured capsules results in extrusion and untwisting of the tubule without undergoing eversion, suggesting that the twisted tubule stores elastic energy to be later transferred into kinetic energy by acting as a spring that is released by relaxation to a cylindrical state^15^. The double-walled structure seen upon relaxation (Fig. 3d, second panel, *arrow*) likely allows the flow of PG matrix from the capsule into the lumen, recharging the forces that push the tubule forward. The process is repeated until the tubule is fully elongated or reaches an obstacle (Supplementary Video 10, 11). In summary, this study demonstrates the operational capability of the nematocyst as a complex and self-assembling biological micromachine. We propose that these ancient and sophisticated organelles represent an ideal model for biologically inspired microscale devices that could be utilized in diverse applications ranging from medical technology to materials science.

## Methods

### Animal husbandry

Animals were raised at 23°C in 12 parts per thousand (ppt) artificial seawater (ASW; Sea Salt; Instant Ocean). Spawning induction and de-jellying and were carried out as previously described^48^. Embryos and polyps were cultured at either room temperature 23C or 25C

### Generation of the *nematogalectin*>*EGFP* transgenic reporter line

The transgenic reporter line was generated by meganuclease mediated insertion^49^ of a plasmid containing EGFP under the control of the nematocyte specific *N. vectensis nematogalectin* promoter.

### *In vivo* labelling of *Nematostella* with (5,6)-tetramethylrohodamine isothiocyanate (TRITC)

Live *Nematostella vectensis* planula (2 dpf) were allowed to react with the amine reactive rhodamine derivative 5/6-tetramethyl-rhodamine-6-isothiocyanate, TRITC (Cayman Chemical, No. 19593) for a short duration (30 min-1 hour). The animals were incubated for 1 hour minutes at a final concentration of 1uM for live imaging of discharge. For fixed specimens, TRITC at a final of 25uM is incubated for 1 hour with 2dpf larvae. The fluorescent dye stained the animals without any apparent toxicity up to 25uM concentration tested in this study (data not shown). Upon incubation, the reactive dye was removed by multiple washes. Animals were transferred to clean dishes in dye free medium for 3 to 5 days until mature nematocytes emerged and the non-specific background fluorescence disappeared substantially. Stock solutions of 25mM TRITC was prepared and stored frozen at −20°C and used without an observable reduction in the chemical reactivity.

### Electron Microscopy

For scanning electron microscopy (SEM) of topography (Fig. 3a), samples were processed according to previous reports^50^. Briefly, samples were fixed in 2.5% glutaraldehyde and 2% paraformaldehyde in 0.1 M NaCacodylate buffer and stained with aqueous tannic acid, osmium tetroxide, thiocarbohydrazide, and then osmium tetroxide again (TOTO). Samples were dehydrated in a graded series of ethanol and critical point dried in a Tousimis Samdri 795 critical point dryer, mounted on stubs, and imaged in a Hitachi TM4000 tabletop SEM at 15kV with BSE detector. For electron microscopy (EM) of internal ultrastructure, samples were fixed as above, with secondary fixation in 1% buffered osmium tetroxide for 1 hour and *en bloc* staining in 0.5% aqueous uranyl acetate carried out overnight at 4°C. A graded series of ethanol was used for dehydration with acetone as a transition solvent and infiltration in Hard Plus resin (Electron Microscopy Sciences). Samples were cured for 48 hours at 60 degrees C and serial sections were cut at 50nm using a Diatome 45-degree ultra-diamond knife or an AT-4 35-degree diamond knife on a Leica UC7 ultramicrotome. Sections were collected on slot grids for STEM imaging and a flat substrate (coverslip or silicon chip) for SEM imaging, and post stained using 4% uranyl acetate in 70% methanol for 4 minutes and Sato’s triple lead stain for 5 minutes. Sections on flat substrate were mounted on stubs, the underside of the coverslip painted with silver paint to mitigate charging, and all were coated with 4 nm carbon in a Leica ACE600 coater. Sections were imaged in a Zeiss Merlin SEM using the aSTEM or 4QBSD detector and Atlas 5 software (Fibics, Inc.). Serial images were aligned and traced for 3D modeling in IMOD, and 3D models rendered in Blender 2.8 (Blender Foundation). Straightening of fully discharged nematocyte (Fig. 2e) was done in Fiji (ImageJ)^60^. Whole nematocyst capsule image composite (fig. 1d) and false coloring and was done in Photoshop 2021 (Adobe, Inc.).

#### Live imaging of nematocyst discharge in live animals

The nematocysts of the TRITC treated animals were discharged by decreasing the pH of the medium using acetic acid. The nematocysts discharge *in vivo* when the medium becomes acidic. Primary polyps were immobilized in glass bottom dishes by sandwiching between a glass slide and the bottom of a glass bottom dish using silicone sealant. The images captured after dropwise addition of glacial acetic acid (37%) to the ASW which triggers capsule discharge when the pH sufficiently decreases in the medium below a certain threshold. Live imaging of the nematocyst maturation and discharge events were recorded with Yokogawa CSU-w1 spinning disc system on a Nikon Ti2 platform with 100X objective.

### Immunofluorescence

TRITC treated primary polyps were fixed in Lavdovski’s fixative (ethanol:formaldehyde:acetic acid:dH_2_O; 50:10:4:36) overnight^20^. The fixative was removed, and the samples were washed 5 times with 1ml PBS pH 7.4 to remove the fixative. Samples were permeabilized with 0.1 % Triton-X100 in PBS, pH 7.4 for 15 min. After several additional washes in PBST (0.1% Tween 20 in PBS, pH 7.4), the polyps were first blocked for 1 hour and incubated over night at 4°C with NvNCol-4 (1:500) in PBST supplemented with 10% goat serum. Minicollagen Ncol4 (rabbit) antibody were kind gifts from Suat Ozbek. The polyps were washed three times in PBST supplemented with 10% goat serum and incubated with ALEXA Fluor 647 (Thermo-Fisher) coupled anti-rabbit secondary antibody (1:500) and WGA-OregonGreen (1:500) overnight in PBST supplemented with 10% goat serum. Thereafter, the polyps were washed several times in PBS and incubated in 90% PBS/Glycerol overnight. The polyps were transferred to a glass slide and mounted on a glass slide with Prolong Glass antifade mounting medium (ThermoFisher). Fluorescence images were acquired using Yokogawa CSU-w1 on a Nikon Ti2 platform. Super-resolution fluorescence confocal images were acquired using the Zeiss LSM780 in Airyscan mode.

### Super-resolution fluorescence confocal imaging

In Fig. 2f, TRITC treated primary polyps were fixed in Lavdovski’s fixative (ethanol:formaldehyde:acetic acid:dH_2_O; 50:10:4:36) overnight^20^. After several washes, the polyps were transferred to PBS/Glycerol (PBS) and incubated overnight. The polyps were transferred to a glass slide and mounted with Prolong Glass antifade mounting medium (ThermoFisher). The polyps were spread onto the glass slides and crushed with the slide cover to disrupt the capsules from the tissue. The fluorescence imaging was performed using the Zeiss LSM780 in Airyscan mode.

### Purification, discharge and staining of TRITC labeled nematocysts

TRITC treated primary polyps were frozen in liquid nitrogen, thawed and macerated manually with a plastic pestle. The samples were suspended in 1 ml Percoll ( 50%, v/v;Sigma) in 300mM sucrose supplemented with 0.01 % Tween20 to prevent adhesion to the microcentrifuge tubes. The tissue is further disrupted by pipetting up and down. The mixture is allowed to settle on ice for 30 minutes and centrifuged for 15 minutes at 3200 rpm. The pellet is washed twice with PBS with 0.01% Tween-20) resuspended in nematocyst discharge buffer (10mM Tris, pH 7.5, 10mM CaCl2). The discharge was initiated by addition of 1mM DTT and incubated for 30 minutes. Upon incubation for 30 minutes, 1ug/ml Wheat germ agglutinin conjugated with OregonGreen (Thermo-Fisher, cat#) was added to the tube and incubated for 1 hour. The stained samples were washed twice with PBST and centrifuged at 3200 rpm for 5 minutes. A loose pellet is seen and suspended in PBST 5ul aliquots were spread onto glass slides and mounted with ProlongGlass (Thermo-Fisher) The images were acquired using Yokogawa CSU-W1 spinning disc system on a Nikon Ti2 platform with 100X objective.

### shRNA knockdown

The short hairpin RNA targeting the putative *Nematostella* spinalin gene (v1g243188) were synthesized by T7 polymerase reaction with the following primers; Forward:TAATACGACTCACTATAGCGGTGGACTCTACTTATTTTCAAGAGAAATAAGTAGAGTCCACCGCTT and Reverse:AAGCGGTGGACTCTACTTATTTCTCTTGAAAATAAGTAGAGTCCACCGCTATAGTGAGTCGTATTA according to the methods described previously^43 44^. Purified shRNA was microinjected into unfertilized eggs at a concentration of 1ug/ul. Following fertilization, the embryos were incubated for 2 days and treated with TRITC.

### Image analysis and mathematical modeling

All image analysis was performed with Fiji^51^. The brightness, contrast and gamma were adjusted manually. The length measurements for the undischarged and discharged capsules and tubules were performed manually using ImageJ freehand manual tracing tool and the results were discussed in supplementary information. The mean signal intensity, area, and length of the objects were acquired in ImageJ by using measurement tool. The results were used in estimates described in supplemental information.

## Acknowledgements

We are grateful to Alejandro Sánchez Alvarado, Jay Unruh, Kausik Si, Whitney Leach, Eric Hill and Subramanian Ramanathan for helpful comments on the manuscript. We are indebted to Molly Simmons for the illustrations of the model. We would like to thank Xia Zhao and Morgan Harwood (Electron Microscopy Core) for the assistance in experiments and the Stowers Reptiles and Aquatics Facility for the animal maintenance. We are thankful to Suat Ozbek for the minicollagen antibodies and Ruohan Zhong for help in generating the transgenic animals. The work was funded by Stowers Institute for Medical Research. Original data of this manuscript can be accessed from the Stowers Original Data Repository.

## Author Contributions

A.K. and M.C.G concepted this study. A.K., B.R. and M.C.G. wrote the manuscript. A.K., S.M.C, M.M. and B.R. performed the experiments and analyzed the data.

## Competing interests

Authors declare no competing interests.

## Materials & Correspondence

The correspondence and materials requests should be addressed to

---

**Supplementary Video 1** | Time lapse of TRITC dye (Magenta) incorporation inside the capsule of a nematocyte expressing EGFP (Green).

**Supplementary Video 2** | Serial SEM of the longitudinal cross section of a basitrichous isorhiza.

**Supplementary Video 3** | 3D reconstruction of the longitudinal cross section of a basitrichous isorhiza showing the connector regions (yellow), the shaft (blue) and a segment of the tubule (magenta) inside the capsule.

**Supplementary Video 4** | Serial SEM of the traverse cross section of a basitrichous isorhiza.

**Supplementary Video 5** | Time lapse of the discharge and eversion of a long basitrichous isorhiza in TRITC labeled polyps. Arrow indicate the capsule embedded inside the tissue.

**Supplementary Video 6** | Discharge events recorded in TRITC labeled polyps in 5msec intervals.

**Supplementary Video 7** | Time lapse of the discharge and partial eversion of a short basitrichous isorhiza in TRITC labeled polyp.

**Supplementary Video 8**| Serial SEM of an everted shaft and the traversing uneverted tubule.

**Supplementary Video 9** | Serial SEM of the longitudinal cross section of an everting thread showing uneverted tubule traversing inside the everted segments.

**Supplementary Video 10**| Time lapse of the movement of a thread possibly blocked by an obstacle showing the bending of the tubule in 180 degree turns.

**Supplementary Video 11** | Time lapse of the discharge and eversion of a short basitrichous isorhiza showing arrested tubule eversion and the forward movement of the uneverted tubule inside the everted fraction.

**Supplementary Information** | Estimates of excess elastic energy released during shaft eversion.

## Notes

### Competing Interest Statement

The authors have declared no competing interest.

## References

1 Tardent, P. The cnidarian cnidocyte, a hightech cellular weaponry. BioEssays 17, 351–362 (1995).

2 Beckmann, A. & Ozbek, S. The nematocyst: a molecular map of the cnidarian stinging organelle. Int J Dev Biol 56, 577–582 (2012).

3 Kass-Simon, G. The behavioral and developmental physiology of nematocysts. Canadian Journal of Zoology-revue Canadienne De Zoologie - CAN J ZOOL 80, 1772–1794 (2002).

4 Watson, G. M. & Hessinger, D. A. Cnidocyte mechanoreceptors are tuned to the movements of swimming prey by chemoreceptors. Science 243, 1589–1591 (1989).

5 Tardent, P. & Holstein, T. Morphology and morphodynamics of the stenotele nematocyst of Hydra attenuata Pall. (Hydrozoa, Cnidaria). Cell Tissue Res 224, 269–290 (1982).

6 Ozbek, S., Balasubramanian, P. G. & Holstein, T. W. Cnidocyst structure and the biomechanics of discharge. Toxicon 54, 1038–1045 (2009).

7 Tardent, P. History and current state of knowledge concerning discharge of cnidae. The biology of nematocysts, 309–332 (1988).

8 Ozbek, S. The cnidarian nematocyst: a miniature extracellular matrix within a secretory vesicle. Protoplasma 248, 635–640 (2011).

9 Lotan, A., Fishman, L., Loya, Y. & Zlotkin, E. Delivery of a nematocyst toxin. Nature 375, 456 (1995).

10 Lotan, A., Fishman, L. & Zlotkin, E. Toxin compartmentation and delivery in the Cnidaria: the nematocyst’s tubule as a multiheaded poisonous arrow. J Exp Zool 275, 444–451 (1996).

11 Holstein, T. & Tardent, P. An ultrahigh-speed analysis of exocytosis: nematocyst discharge. Science 223, 830–833 (1984).

12 Nuchter, T., Benoit, M., Engel, U., Ozbek, S. & Holstein, T. W. Nanosecond-scale kinetics of nematocyst discharge. Curr Biol 16, R316–318 (2006).

13 Weber, J. Nematocysts (stinging capsules of Cnidaria) as Donnan-potential-dominated osmotic systems. Eur J Biochem 184, 465–476 (1989).

14 Weber, J. A novel kind of polyanions as principal components of cnidarian nematocysts. Comparative Biochemistry and Physiology Part A: Physiology 98, 285–291 (1991).

15 Carre, D. Hypotèse sur le mécanisme de l’évagination du filament urticant des cnidocystes Eur J Cell Biol 20, 265–271 (1980).

16 Godknecht, A. & Tardent, P. Discharge and mode of action of the tentacular nematocysts of Anemonia sulcata (Anthozoa: Cnidaria). Marine Biology 100, 83–92 (1988).

17 Weill, R. Contribution a l’étude des cnidaires et de leurs nématocystes. (Laboratoire d’Èvolution des Ítres organisÈs : Les presses universitaires de France, 1934).

18 Östman, C. A guideline to nematocyst nomenclature and classification, and some notes on the systematic value of nematocysts. Scientia Marina 64, 31–46 (2000).

19 Fautin, D. G. Structural diversity, systematics, and evolution of cnidae. Toxicon 54, 1054–1064 (2009).

20 Zenkert, C., Takahashi, T., Diesner, M. O. & Ozbek, S. Morphological and molecular analysis of the Nematostella vectensis cnidom. PLoS One 6, e22725 (2011).

21 Babonis, L. S. & Martindale, M. Q. PaxA, but not PaxC, is required for cnidocyte development in the sea anemone Nematostella vectensis. Evodevo 8, 14 (2017).

22 Reft, A. J., Westfall, J. A. & Fautin, D. G. Formation of the apical flaps in nematocysts of sea anemones (cnidaria: actiniaria). Biol Bull 217, 25–34 (2009).

23 Watson, G. M. & Mariscal, R. N. Ultrastructure of nematocyst discharge in catch tentacles of the sea anemone Haliplanella luciae (Cnidaria: Anthozoa). Tissue Cell 17, 199–213 (1985).

24 Park, S. et al. The nematocyst’s sting is driven by the tubule moving front. J R Soc Interface 14 (2017).

25 Aerne, B., Stidwill, R. & Tardent, P. Nematocyst discharge in Hydra does not require the presence of nerve cells. Journal of Experimental Zoology - J EXP ZOOL 258, 137–141 (1991).

26 Watson, G. M. & Hessinger, D. A. Cnidocytes and adjacent supporting cells form receptor-effector complexes in anemone tentacles. Tissue Cell 21, 17–24 (1989).

27 Westfall, J. A., Landers, D. D. & McCallum, J. D. Different nematocytes have different synapses in the sea anemone Aiptasia pallida (Cnidaria, Anthozoa). Journal of Morphology 238, 53–62 (1998).

28 Thorington, G. U. & Hessinger, D. A. Efferent Mechanisms of Discharging Cnidae: II. A Nematocyst Release Response in the Sea Anemone Tentacle. Biol Bull 195, 145–155 (1998).

29 Weir, K., Dupre, C., van Giesen, L., Lee, A. S. & Bellono, N. W. A molecular filter for the cnidarian stinging response. Elife 9 (2020).

30 Holstein, T. W. et al. Fibrous mini-collagens in hydra nematocysts. Science 265, 402–404 (1994).

31 David, C. et al. Evolution of complex structures: Minicollagens shape the cnidarian nematocyst. Trends in genetics : TIG 24, 431–438 (2008).

32 Adamczyk, P. et al. Minicollagen-15, a novel minicollagen isolated from Hydra, forms tubule structures in nematocysts. J Mol Biol 376, 1008–1020 (2008).

33 Sunagar, K. et al. Cell type-specific expression profiling unravels the development and evolution of stinging cells in sea anemone. BMC Biol 16, 108 (2018).

34 Weber, J. Novel tools for the study of development, migration and turnover of nematocytes (cnidarian stinging cells). J Cell Sci 108 ( Pt 1), 403–412 (1995).

35 Krohne, G. Organelle survival in a foreign organism: Hydra nematocysts in the flatworm Microstomum lineare. Eur J Cell Biol 97, 289–299 (2018).

36 Engel, U. et al. Nowa, a novel protein with minicollagen Cys-rich domains, is involved in nematocyst formation in Hydra. J Cell Sci 115, 3923–3934 (2002).

37 Yamada, S., Morimoto, H., Fujisawa, T. & Sugahara, K. Glycosaminoglycans in Hydra magnipapillata (Hydrozoa, Cnidaria): demonstration of chondroitin in the developing nematocyst, the sting organelle, and structural characterization of glycosaminoglycans. Glycobiology 17, 886–894 (2007).

38 Yamada, S., Sugahara, K. & Ozbek, S. Evolution of glycosaminoglycans: Comparative biochemical study. Commun Integr Biol 4, 150–158 (2011).

39 Hwang, J. S. et al. Nematogalectin, a nematocyst protein with GlyXY and galectin domains, demonstrates nematocyte-specific alternative splicing in Hydra. Proc Natl Acad Sci U S A 107, 18539–18544 (2010).

40 Adamczyk, P. et al. A non-sulfated chondroitin stabilizes membrane tubulation in cnidarian organelles. J Biol Chem 285, 25613–25623 (2010).

41 Itakura, Y., Nakamura-Tsuruta, S., Kominami, J., Tateno, H. & Hirabayashi, J. Sugar-Binding Profiles of Chitin-Binding Lectins from the Hevein Family: A Comprehensive Study. Int J Mol Sci 18, 1160 (2017).

42 Skaer, R. J. & Picken, L. E. The pleated surface of the undischarged thread of a nematocyst and its simulation by models. J Exp Biol 45, 173–176 (1966).

43 He, S. et al. An axial Hox code controls tissue segmentation and body patterning in Nematostella vectensis. Science 361, 1377–1380 (2018).

44 Karabulut, A., He, S., Chen, C. Y., McKinney, S. A. & Gibson, M. C. Electroporation of short hairpin RNAs for rapid and efficient gene knockdown in the starlet sea anemone, Nematostella vectensis. Dev Biol 448, 7–15 (2019).

45 Sebe-Pedros, A. et al. Cnidarian Cell Type Diversity and Regulation Revealed by Whole-Organism Single-Cell RNA-Seq. Cell 173, 1520–1534 e1520 (2018).

46 Hellstern, S. et al. Structure/function analysis of spinalin, a spine protein of Hydra nematocysts. FEBS J 273, 3230–3237 (2006).

47 Koch, A. W. et al. Spinalin, a new glycine- and histidine-rich protein in spines of Hydra nematocysts. J Cell Sci 111 ( Pt 11), 1545–1554 (1998).

48 Genikhovich, G. & Technau, U. Induction of spawning in the starlet sea anemone Nematostella vectensis, in vitro fertilization of gametes, and dejellying of zygotes. Cold Spring Harb Protoc 2009, pdb prot5281 (2009).

49 Renfer, E. & Technau, U. Meganuclease-assisted generation of stable transgenics in the sea anemone Nematostella vectensis. Nat Protoc 12, 1844–1854 (2017).

50 Jongebloed, W. L., Stokroos, I., Van der Want, J. J. & Kalicharan, D. Non-coating fixation techniques or redundancy of conductive coating, low kV FE-SEM operation and combined SEM/TEM of biological tissues. J Microsc 193, 158–170 (1999).

51 Schindelin, J. et al. Fiji: an open-source platform for biological-image analysis. Nature Methods 9, 676–682 (2012).

