## Supplementary Information for "The architecture and operating mechanism of a cnidarian stinging organelle"

### 1 Helix elastic energy

Consider a filament of radius $r$ bent into a helix (no twisting assumed to simplify calculations) with radius $R$ and pitch $c$ defined by the parametric equation

$$x = R\cos t, \quad y = R\sin t, \quad z = ct. \tag{1}$$

The curvature of such a helix is $\kappa = R/(R^2 + c^2)$. The bending energy $e$ per unit length is given by [1]

$$e = \pi Y_0 \kappa^2 r^4/8, \tag{2}$$

where $Y_0$ stands for the filament Young modulus. For tightly coiled structure (its segment is shown in Fig.1) we obtain that $2\pi c_c = 6r$ giving $c_c = 3r/\pi \approx r$. The radius of this coil $R_c = r$ and the curvature $\kappa_c = 1/(2r)$. Thus we find the bending energy $E_c$ of the filament of the length $L$

$$E_c = \pi Y_0 L r^2/32 \approx Y_0 L r^2/10. \tag{3}$$

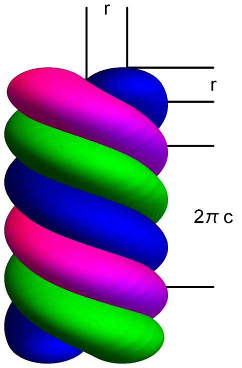


Figure 1: A segment of tightly coiled shaft before eversion. The diameter of the shaft is equal to double diameter of the filament while the pitch $c$ satisfies the condition $2\pi c = 6r$.

Consider an uncoiled helix with the increased radius $R_{uc} = kR_c = kr,\ k > 1$ and pitch $c_{uc} = mc_c = mr,\ m > 1$. Thus the curvature of this helix reads $\kappa_{uc} = k/((k^2+m^2)r) = 2k/(k^2+m^2)\kappa_c$. This leads to the expression for the bending energy

$$E_{uc} = 4k^2 E_c/(k^2+m^2)^2. \tag{4}$$

The measurements of the geometric parameters of the coiled and uncoiled helices produce the following values (in $\mu$m)

$$R_c = r = 0.12,\ R_{uc} = 0.6,\ c_c = r = 0.12,\ c_{uc} = 1.2,$$

leading to $k = 5$ and $m = 10$. Thus the energy stored in the uncoiled structure is less than 1% of that stored in the coiled structure. Similarly one can neglect the energy spent into torsion of the tubule attached to the lower tip of the uncoiling shaft. This estimate means that nearly all bending energy stored in the coiled structure $E_b = 3E_c$ can be converted into kinetic energy $E_k = 3E_c$ of the tubule motion.

#### 2 Minicollagen fibers material parameters

The staining of the shaft filaments shows that they are made of minicollagens. The structural analysis of these proteins show that they have CRD regions, polypropiline segment(s) and more flexible collagen motif. Similar structure was also observed in the CPP-1 molecule found in *Hydra* nematocyst that has Young modulus of $Y = 7.8 \pm 8.0$ MPa in the bulk [2]. Collagen fibers demostrate Young modulus values order of few hundreds MPa found using different methods [3, 4]. It is convenient to use $Y$ as an estimate for $Y_0$. This gives us an estimate for $E_k$ for the filaments of the length $L = 20$ $\mu$m

$$E_k = 3E_c = 0.3YLr^2 = 1\ pJ.$$

#### 3 Tubule velocity and viscous drug estimates

The tubule is pulled out of the capsule using the elastic energy released during the shaft eversion. It covers the distance $\mathcal{L}$ moving in the prey tissue with the average velocity is $u = 1.5\ mm/s$. This value allows to estimate the Reynolds number $Re$ of such motion using the definition $Re = 2\rho_w ur/\mu_v$, where $\rho_w = 10^3 kg/m^3$ and $\mu_v = 9{\cdot}10^{-4}\ Pa{\cdot}s$ denote water density and dynamic viscosity respectively. We find $Re = 0.002$ that implies that one can neglect the inertial effects which means that only viscous drag matters for the tubule motion.

First note that in this slow motion liquid inside the tubule is stationary with respect to the tubule wall so that the drag can be approximated by the one of the torus moving with constant velocity $u$ along its main axis. If we denote $w$ the tubule wall thickness the major $\mathcal{R}$ and minor $\rho$ axes of the torus read

$$\mathcal{R} = r - w/2, \qquad \rho = w/2.$$

Then the drag force $F_d$ and corresponding work $A_d$ are found as

$$F_d = 6\pi\mu_v u\rho f, \quad A_d = F_d\mathcal{L} = 6\pi\mu_v u\rho\mathcal{L}f.$$

Here $f$ denotes a nondimensional function of the ratio $\mathcal{R}/\rho$ in the range of few units. We choose $f = 3$ and find with $w = 0.1\ \mu m$ the estimate for the drag work

$$A_d = 0.25 \cdot 10^{-15}\ J = 0.25\ fJ.$$

We conclude that this work is several orders of magnitude smaller that the elastic energy stored in the coiled shaft.

#### 4 Energy cost of tubule eversion

Using the elastic energy of the everting shaft the tubule also can evert. To estimate the energy cost of this process one has to consider a tubule as an elastic cylinder of radius $r$ and length $\mathcal{L}$ everting inside out. With some assumptions about the tubule elastic properties such a problem can have an exact solution for the evertion work $A_e$ that reads

$$A_e = \pi\mu\mathcal{L}r^2(1 - \delta^2),$$

where $\mu$ is the is the shear modulus of the tubule material and $\delta$ is the ratio of the inner tubule radius to the outer one. Taking $\delta = 0.8$ and using shear modulus of the collagen $\mu = 35\ MPa$ we find

$$A_e = 0.125 \cdot 10^{-9}\ J = 0.125\ nJ.$$

We observe that this energy is three orders of magnitude larger than that of provided by the elastic energy of the shaft so that tubule eversion is possible if other sources of energy are included into consideration.

#### References


[1] Landau LD, Lifshitz EM, *Theory of Elasticity*, Pergamon Press, 1970.

[2] Bentele T, Amadei F *et al*, New Class of Crosslinker-Free Nanofiber Biomaterials from Hydra Nematocyst Proteins, Sci.Reports, 2019, 9:19116.

[3] Dutov P, Antipova O *et al*, Measurement of ElasticModulus of Collagen Type I Single Fiber, PLoS ONE, 2016, 11(1): e0145711.

[4] Van der Rijt JAJ, van der Werf KO *et al*, Micromechanical testing of individual collagen fibrils, Macromol Biosci., 2006; 6: 697702.

[5] Weber J, Klug M, and Tardent P, Some physical and chemical properties of purified nematocysts of Hydra attenuata Pall. (Hydrozoa, Cnidaria), Comp. Biochem. Physiol. 1987; 88B: 855862.
